# Could microalgae serve as an alternative to replace unsustainable palm oil?

**DOI:** 10.1101/2025.04.28.651004

**Authors:** Karolína Štěrbová, Kateřina Bišová, João Artur Câmara Manoel, Jiří Masojídek

## Abstract

Palm oil is the world’s most widely used vegetable oil with a significant impact on the environment. Microalgae produce a variety of fatty acids whose composition is influenced by growth conditions. We selected a set of ten algal strains that naturally produce oils similar in composition to palm oil, and we analyzed the effects of varying light intensity on their growth and fatty acid production and composition while maintaining temperatureThe optimum irradiance in terms of biomass accumulation was determined to be 400 µmol photons m^−2^ s^−1^ for most of the microalgae, except for *C. moewusii* CCALA 242 and CCALA 243, where the optimum illuminance was determined to be 100 and 50 µmol photons m^−2^ s^−1^, respectively. The growth of *Scenedesmus* and *Desmodesmus* strains improved with increasing light intensity and mostly reached the growth rates approximately twice as high as the *C. moewusii* strains at higher light intensities. In the presence of NaCl, the growth of *C. moewusii* CCALA 244 and *D. subspicatus* CCALA 467 was similar or better than under the non-stress conditions. In contrast, the growth of *D. communis* CCALA 463 was reduced and biomass productivity decreased. In *C. moewusii* CCALA 244, salinity improved total fatty acid production 5-fold and had a particular effect on the production of palmitic acid, stearic acid, oleic acid and linolenic acid. The lab-scale growth could be successfully recapitulated in large-scale photobioreactor, although the total fatty acid productivity was lower. Interestingly, *D. communis* CCALA 463 showed a 3.5-fold improvement in growth in the large-scale photobioreactor as well as improved total fatty acid productivity. Eight of the ten microalgal strains used were capable of heterotrophic growth. Although specific growth rates were higher in heterotrophy, biomass productivity and total fatty acid productivity were lower than in the case of high light intensities of the autotrophic treatment.

## Introduction

Palm oil is the most widely used vegetable oil worldwide accounting for 37% of annual production with a large range of industrial applications while oil production from soybean, rapeseed and sunflower make up 28, 12 and 10%, rspectively [1]. Nevertheless, about half of all consumer products sold in supermarkets are made of palm oil [2]. The world’s leading producers of palm oil are Indonesia and Malaysia, which make up more than 85% of the global palm oil supply [3,4]. By 2050 the overal demand for palm oil is predicted to reach 240 million tonnes [5].

Despite being inexpensive, palm oil is known for its significant environmental impact. Deforestation and the use of palm trees triggered by oil production are serious issues linked to global warming. Thus, there is an urgent need for researach on alternatives [6,7]. The principal constituent of palm oil is palmitic acid (PA) accounting for almost half of the amount (44%). Then, it contains oleic acid (OA; 39%), linoleic acid (LA; 10%) and the rest are small amounts represented by stearic, myrystic, linolenic, lauric and arachidic acids [8]. Microalgae produce a variety of saturated (SFA), unsaturated (UFA) and polyunsaturated (PUFA) fatty acis and the extracted oil has a similar chemical composition to the palm oil making them an alternative source. Moreover, their cultivation is environmentally sustainable taking less space to grow with a higher yield than palm trees. This makes them a promising alternative for research to replace or at least reduce the demand for palm oil [6]. Many recent studies have revealed the health potentials associated with the monounsaturated fatty acids from microalgae [9].

As the lipid production and fatty acid profile are related to the cultivation conditions, there is a need to find the most suitable cultivation conditions for individual microalgae species producing an adequate amount of targeted FAs. The fatty acids are important constituents of microalgae biomass and typically count for 5 to 60% of cell dry weight. The compounds and amounts are species-specific [10]. Palmitic acid (PA), the most abundant FA in palm oil, can be found in several phyla such as Chlorophyta, Rhodophyta, Haptophyta, Cryophyta, Dinophyta and Bacillariophyta if grown under nutrient-replete conditions [11]. As mentioned above, cultivation variables such as light intensity, temperature and nutrient limitation influence the FA profile and hence, the quality and quantity of lipids produced [12]. As we deal with photosynthetic photoautotrophs, irradiance is one of the most important variable for microalgae growth in general [13,14] and the suitable level depends on species, varying between tens to hundreds micromols photons per m^−2^ s^−1^ [10,14]. Higher light intensity increases lipid content [15,16] and together with exposure duration is related to variations of the contents SFAs, MUFAs and PUFAs [10,17]. In most cases, microalgae are cultivated in photoautotrophic conditions, but several species can also grow under heterotrophic conditions in the dark in the presence of organic carbon sources (e.g. glucose, fructose, sucrose, lactose, galactose, acetic acid [18,19]. In adition, some studies have reported the improvement of lipid production under mixotrophic cultivation although not all microalgae species are suitable for heterotrophic or mixotrophic cultivation [20].

Irradiance is not the only factor playing an important role in microalgae growth and changes in FA profiles in a wide range of microalgae species. The response of chemical composition to high and low temperatures is species dependent. The suitable growth temperature for each species is related to the temperature at the collection location and it ranges between 21 and 29°C (except Cryophyta where T_opt_=16°C). The changes in FA profile of microalgae to high and low growth temperatures vary from species to species [16]. The total content of lipids is usually enhanced with increasing temperature [21]. Specifically, microalgae tend to produce and accumulate SFAs as the temperature increases; at low temperatures, microalgae usually accumulate UFAs [16].

Another way to modulate the content of lipids is nutrient deprivation. Nitrogen (N) is a major component in many macromolecules like chlorophylls, proteins, and DNA [22] and it is present in microalgae cell in the amount between 1-10% of cell dry weight [23]. At an industrial scale, it is added to the cultivation medium as ammonium, urea or nitrate and its deficiency causes a decrease in growth and induces the production of lipids [24] even though their production rate is decreased due to a slowdown in microalgae growth [25]. Under N depletion, microalgae grow in a medium lacking N source, while under N limitation, there is a constant but insufficient amount available and cells are still growing at a reduced rate given by limiting nitrogen uptake [22]. Nitrogen limitation is one of the most cost-effective and easily adjustable conditions for enhancing lipid accumulation [16].

Most studies have focused on the effects of light, temperature, and N-deprivation on biochemical composition. However, another abiotic factor affecting lipid content is salinity. Usually, an increase in medium salinity causes an increase in lipid content [26–28]; however, an excess of NaCl in the medium reduces biomass and lipid productivity [29].

In this study, several Chlorophyta species were grown in photoautotrophic as well as heterotrophic cultivation regimes in different laboratory bioreactors and the quality and quantity of individual FAs reflecting the profile of palm oil were studied at various cultivation conditions.

## Materials and methods

### Organisms and culture maintenance

Ten different microalgae belonging to genera *Chlamydomonas, Scenedesmus* and *Desmodesmus* (class Chlorophyceae; obtained from the Culture Collection of Autotrophic Organisms, Třeboň, Czech Republic) were selected based on the highest amounts of PA and OA, the two most abundant FAs in palm oil [30]. Three strains of microalgae species *Chlamydomonas moewusii* (CCALA 242, CCALA 243 and CCALA 244), three strains of species *Scenedesmus obliquus* (CCALA 453, CCALA 455 and CCALA 456), two strains of species *Desmodesmus communis* (CCALA 463 and CCALA 464), and strain of species *Scenedesmus subspicatus* (CCALA 688) and *Desmodesmus subspicatus* (CCALA 467) were used.

In the case of photoautotrophic cultivation, the cultures were initially grown in the BG-11 medium [31,32] in 250 mL Erlenmayer flasks placed on the shaker in the laboratory at 22-25°C and illuminated by continuous light of about 50 µmol photons m^−2^ s^−1^.

For heterotrophic cultivation, the strains were maintained on agar solidified ½ ŠS medium (Hlavová et al., 2016) by subculturing every three weeks. The freshly streaked cultures were grown on a light shelf at an incident light intensity 100 µmol m^−2^ s^−1^ photosynthetically active radiation at 22-25 °C for about a week and then stored in dark at 15 °C.

### Photoautotrophic cultivation

The light optimization (Trial 1) of all microalgae was performed in 100 mL glass columns with a light path of 25 mm. Each column was inoculated with a culture to reach the initial average optical density of about OD_750_=0.2. The columns were submerged in a temperature-controlled water bath set to the growth optimum of 30°C which is common for green microalgae [33]. The columns were mixed by bubbling air+1% CO_2_ (v/v) with a flow rate of 0.1 L min^−1^ and exposed to four light intensities 50, 100, 200 and 400 µmol photons m^−2^ s^−1^ to find the suitable value of irradiation providing the desired biochemical response. The experiment lasted for 10 days. The suitable light intensity and total content of FAs as well as the content of individual FAs were determined. From the total number of ten microalgae, three (*C. moewusii* CCALA 244, *D. communis* CCALA 463 and *D. subspicatus* CCALA 467) were selected for further investigation based on the highest amount of PA and OA.

In the second photoautotrophic trial (Trial 2), the growth and content of FAs of selected microalgae strains based on the criterion described earlier were examined using the same apparatus and columns as in Trial 1 but under stress conditions characterized by sub-optimal temperature of 25°C to increase UFAs [34] or using the synergism of both sub-optimal temperature and addition of NaCl, where a concentration of 16 g L^−1^ was selected as suitable for targeted SFA accumulation [35,36]. The suitable light intensity of 100, 200 and 400 µmol photons m^−2^ s^−1^ was set for *D. communis* CCALA 463*, C. moewusii* CCALA 244 and *D. subspicatus* CCALA 467, respectively based on the previous trial. The initial biomass density was set between 0.2 and 0.3 g dry weight (DW) L^−1^. The growth and biochemical composition in terms of FA composition were followed for 10 days and the impact of stress conditions was determined.

In the photoautotrophic Trial 3, the growth and FA content of three microalgae already used in Trial 2 were verified in larger-scale cultivation. Three microalgae, *D. communis* CCALA 463*, C. moewusii* CCALA 244 and *D. subspicatus* CCALA 467, were separately grown in an annular column photobioreactor (AC-PBR) with a volume of 30 L [37,38]. The temperature was set to 25°C and the initial incident light intensity to 200 µmol photons m^−2^ s^−1^ which was gradually increased to 1600 µmol photons m^−2^ s^−1^ for each individual microalgae to maintain the same conditions. The light pattern is described in Table 1. The culture was mixed by bubbling air+1% CO_2_ (v/v) with a flow rate of 2 L min^−1^. The initial biomass density was set between 0.4 and 0.7 g L^−1^ and the individual trials lasted for 21 days.

**Table 1.** Light pattern set during the trial in AC-PBR measured by LI-COR quantum sensor inside the culture.

| Day of cultivation | Light intensity [$\mu\text{mol photons m}^{-2} \text{s}^{-1}$] |
| --- | --- |
| 1-2 | 200 |
| 2-3 | 300 |
| 3-4 | 400 |
| 4-5 | 500 |
| 5-6 | 600 |
| 6-7 | 800 |
| 7-8 | 1000 |
| 8-9 | 1200 |
| 9-11 | 1400 |
| 11-21 | 1600 |

All cultivation trials were carried out in triplicates.

### Heterotrophic cultivation

For the experiments, the cultures were inoculated directly from the plates into 300 ml of ½ ŠS medium and grown autotrophically in glass cylinders (inner diameter 36 mm, height 500 mm, volume of suspension 300 mL) at countinuous light of incident light intensity 500 µmol m^−2^ s^−1^, at 30 °C and were aerated with 2 % CO_2_ (v/v) in air. The cultures were grown until reaching optical density at 750 (OD_750_) above 0.3; then they were diluted to approximately 10^6^ cell/ml in ½ ŠS medium containing either 1 or 2 % glucose as carbon source and placed into an RTS-8 multi-channel bioreactor (Biosan, Latvia) with the following settings: culture volume of 40 mL, temperature of 30 °C, and agitation 2,000 rpm. The growth of the culture was monitored as changes in optical density at 660 nm. The composition of the biomass was analyzed at the end of the cultivation. All cultivation trials were carried out in triplicates.

## Analytical measurements

### Biomass density

The biomass density of the culture was expressed as dry weight (DW) and the measurement was performed by filtering culture samples on pre-weighed glass microfiber filters (GC-50) as described previously [33]. The filters with the biomass were washed twice with deionized water and dried in an oven at 105 °C for 8 h; then they were weighed (precision of ± 0.01 mg) and the biomass amount (g DW per litre) was calculated. For the DW determination of microalgae cultivated in the medium supplemented by NaCl, the culture suspension was filtered, washed twice with 0.05 M ammonium formate [39], and dried as described previously.

The biomass concentration was also estimated by measuring the optical density (OD_750_) as described previously [40]. A high-resolution spectrophotometer (UV 2600 UV-VIS, Shimadzu, Japan) was used for the measurement.

The specific growth rate µ = (ln DW_2_ – ln DW_1_)/ t_2_ – t_1_ (d^−1^) was calculated throughout the exponential phase and the volumetric as well as areal productivity was calculated according to the following equations: P_V_=((DW_2_-DW_1_)/t_2_ – t_1_) (g DW L^−1^ d^−1^); P_A_= (Pv×V)/S (g DW m^−2^ d^−1^), where V is the volume of culture (L) and S is the illuminated surface area (m^2^).

### Cell counting

Samples for cell counting were prepared according to the protocol described previously [38]. The volume of 500 µL of microalgae culture was fixed with 50 µL of 25% glutaraldehyde and stored at 4 °C until analysis. The cells were counted using a 50 µm aperture in a Beckman Coulter Multisizer^TM^ 4 Coulter Counter^®^.

### Nutrient analysis

The content of the major limiting nutrient, nitrate (NO_3_^−^) in the cultivation medium was determined as described earlier [41]. Samples (5 mL) taken from microalgae cultures were centrifuged at 2,300 × *g* for 10 min and the supernatant was collected and stored at –20 °C before analysis. The samples were diluted five to ten times according the expected NO_3_^−^ concentration with deionized water and the nitrate analysis was performed using an ion chromatography system (ICS-90, Dionex fitted with an AS22-Fast 4×150 mm, Dionex IonPac^TM^ column). A solution of known NaNO_3_ concentration was used as the standard to construct the calibration curve. Nitrate concentrations were calculated (in mg L^−1^) and expressed as a percentage of the initial concentration (% of initial).

### Analysis of fatty acids

The identification as well as quantification of FAs were performed in the biomass samples taken at the end (D10) of the trial. The analysis was performed according to the protocol described previously [37,38]. The amount corresponding to 5-10 mg of lyophilized microalgae biomass was transferred to a breaking vial; 0.4 mL of zirconium/silicon beads (⌀ 0.1 mm), 1 mL mixture of 3 M hydrochloric acid in methanol and 50 µg of internal standard (C15:0) were added. Then, the microalgae cells were disintegrated using 5 consecutive 30 s cycles, using a bead beater (Mini-Beadbeater-16, BioSpec Products, USA). After the disintegration, the samples were cooled down on ice. Then, the content of the breaking vial was transferred into a screw-top test tube and the breaking vial was washed twice with 1 mL of methanol. The sample was sonicated for 15 min (Kraintek 6, Czech Republic) and then, the reaction mixture was heated to 90°C for 1.5 h in a thermoblock. The tubes were then removed from the thermoblock, and cooled to laboratory temperature, and 2 mL of hexane and 2 mL of 1 M NaCl were added. The mixture was vortexed for 10 s and centrifuged at 900 × *g* at 4°C for 10 min (Eppendorf centrifuge 5804 R). The upper organic phase was separated and analysed.

The separation of methyl esters of individual fatty acids (FAMEs) was performed using a Thermo Trace 1300 gas chromatography system with a TR-FAME column (60 m × 0.32 mm, df 0.25 μm) and helium was used as a carrier gas at a pressure of 200 kPa. The temperature ramp was as follows: the starting value was 140 °C which was increased to 240 °C at a rate of 4.5 °C per minute and then maintained at 240 °C for 10 min. The injector was kept at 260 °C and the detector at 250°C. The retention times of FAMEs were compared to known standards from menhaden fish oil (Supelco® 37 Component FAME Mix; PUFA No.3 Supelco). The amount of individual fatty acids was evaluated using internal standards with the known amount of glycerol tripentadecanoate (C15:0) and calculated by multiplying the integrated peak areas by the correction factors of the FID response. In total 13 different FAs (C12:0, C14:0, C16:0, C16:1n7, C16:2, C16:3, C18:0, C18:1n7, C18:1n9, C18:2n6, C18:3n3, C18:3n6 and C20:0) were analysed but only those present at concentrations higher than 10% of TFA were considered.

### Statistical analysis

All measurements were performed in triplicate (*n* = 3); the means and standard deviations (SD) are reported in the figures. Sigma Plot 11.0 was used to determine significant differences between treatments. One-way analysis of variance (ANOVA) and the Holm-Sidac test were conducted for comparison of variables in the trials. P values lower than 0.05 were considered to be significantly different. In graphs, the mean values designated by the same letter did not differ from each other.

## Results

In the first series of experiments (Trial 1), the cultures were illuminated by different light intensities (50, 100, 20 and 400 µmol photons m^−2^ s^−1^) to determine the suitable irradiance of individual microalgae and subsequently the optimum harvesting point to obtain biomass with highest PA and OA concentration. In the second series of experiments (Trial 2), only three of ten microalgae, *D. subspicatus* CCALA 467*, D. communis* CCALA 463 and *C. moewusii* CCALA 244 based on the highest amount of PA and OA, were selected for further investigation. These microalgae were cultivated at the suitable light intensity determined in Trial 1 under unfavourable conditions (lower temperature and salt stress; see detailed information in Materials and methods) to induce the prodution of desirable compounds. The regimes set up in Trial 2 were verified on a larger scale in 30 L AC-PRB (Trial 3). To analyze if the ten algal species can grow hereterotrophically, they were all tested on medium containing glucose. All of them, with the exception of *C. moewusii* CCALA 243 and CCALA 242, grew heteretrophically and thus auto– and heterotrophic growth can be compared.

## Trial 1: Determination of suitable light intensity

In Trial 1, the suitable light intensity of all microalgae was determined based on the accumulation of the biomass and therefore the calculated maximum growth rate. All cultures grew well (Fig. 1) except *Chlamydomonas* species (CCALA 242, CCALA 243 and CCALA 244) as considerable sedimentation of the cells was observed (mostly in *C. moewusii* CCLA 243). In most cases, the value of 400 μmol photons·m^−2^ s^−1^ was determined as suitable except for two *C. moewusii* strains (CCALA 242 and CCALA 243) species in which the highest biomass accumulation was observed at 100 and 50 μmol photons·m^−2^ s^−1^, respectively.

**Fig. 1.**
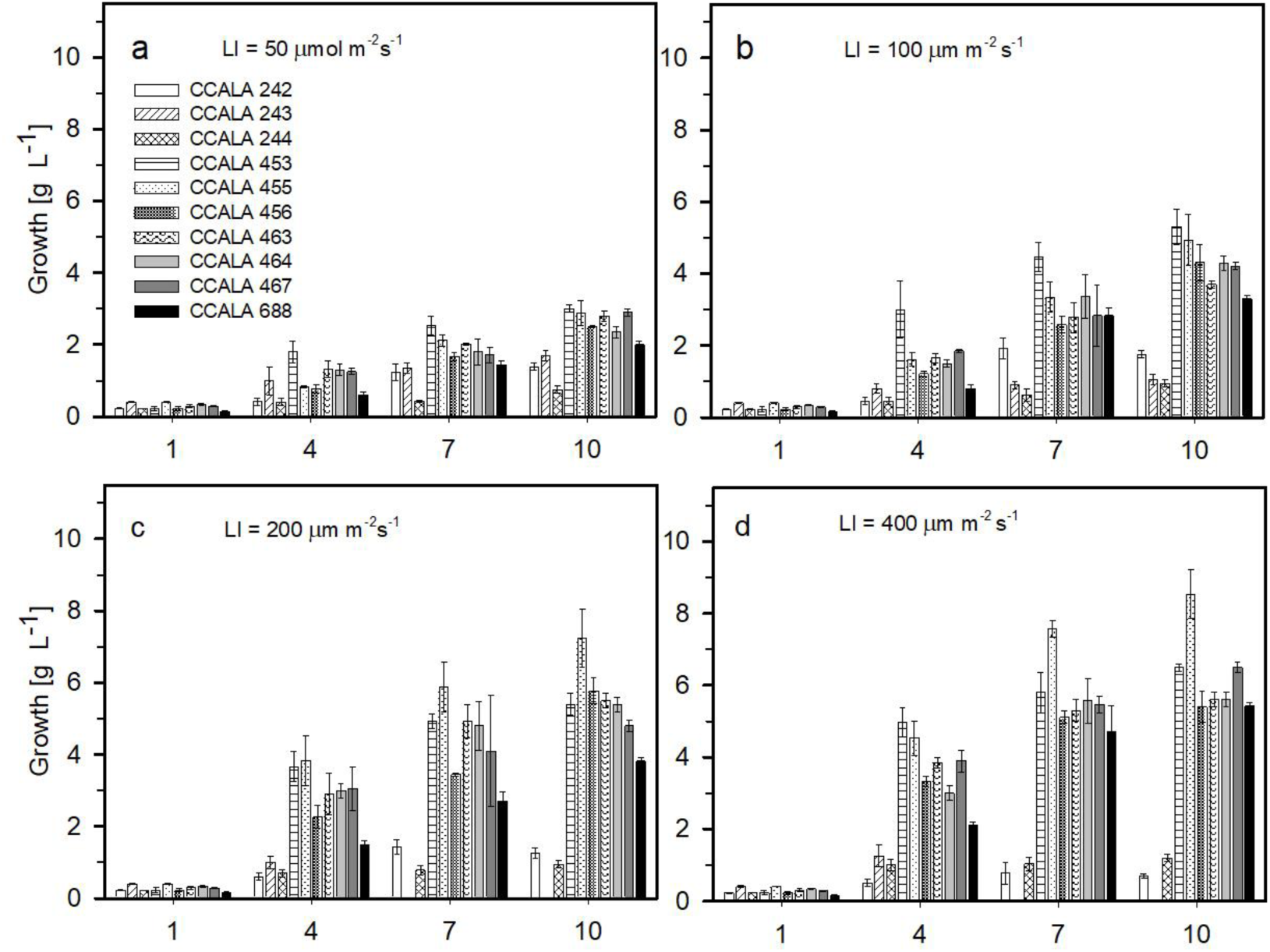
Growth of selected microalgae (*Scenedesmus obliquus* strains CCALA 453, CCALA 455 and CCALA 456, *Chlamydomonas moewusii* strains CCALA 242, CCALA 243 and CCALA 244, *Desmodesmus communis* strains CCALA 463 and CCALA 464, *Scenedesmus subspicatus* CCALA 688, *Desmodesmus subspicatus* CCALA 467) followed at different light intensities for 10 days. The values are presented as a mean (n = 3) ± SE and those designated by the same letter did not differ significantly from each other.

The highest growth rate was determined for *S. subspicatus* CCALA 688 reaching the value of µ=0.58 d^−1^. On the other hand, the highest biomass density of 8.53 ± 0.68 g L^−1^ was reached in *S. obliquus* CCALA 455 (Fig. 1d) though the growth rate was lower µ=0.49 d^−1^ (Fig. 2).

**Fig. 2.**
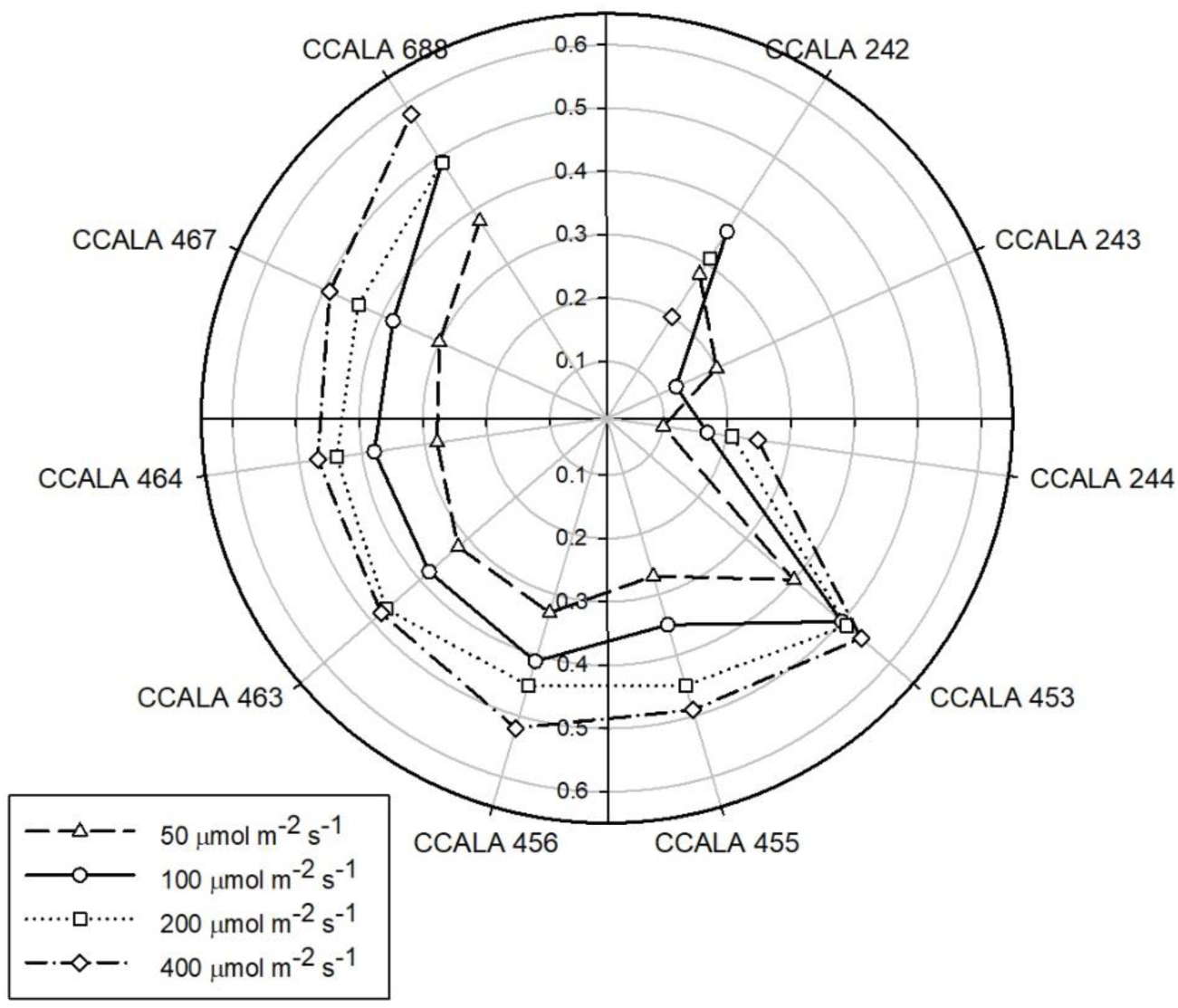
Values of the specific growth rate of selected microalgae strains (see legend of Fig. 1) when exposed to various irradiance levels of 50, 100, 200 and 400 μmol photons m^−2^ s^−1^.

In general, the cultures of the *C. moewusii* strains did not grow well. On the other hand, the culture of *S. obliquus* CCALA 455 grew to high maximum densities, no limitation by nutrient was found. This statement is supported by the analysis of nitrate, a form of nitrogen source as a major macronutrient. From the initial concentration of 1.36 ± 0.03 g L^−1^ of nitrate, even at the highest light intensity of 400 μmol photons m^−2^ s^−1^ with the highest growth rate, the concentration at the end of the trial was about

0.1 g L^−1^ which is still 6.5% of the initial amount (Table 2). On the other hand, a nutrient limitation was observed in *D. commmunis* CCALA 464 culture grown at 400 μmol photons m^−2^ s^−1^ as there was a decline in growth at the end of the trial as no nitrate was analysed. Limitation by light is excluded because at 200 μmol photons m^−2^ s^−1^ the culture growth continued at a comparable density as the culture grown at 400 μmol photons m^−2^ s^−1^. The light limitation was also observed during the cultivation of *S. subspicatus* CCALA 688 at 50 μmol photons m^−2^ s^−1^. The stationary phase was reached after 5 days of cultivation and on the last day of the trial, there was still sufficient amount of nitrate in the cultivation medium coresponding to 29% of the initial concentration (Table 2).

**Table 2.** The concentration of NO^−^ measured at the end of the trial with selected microalgae (see legend of Fig. 1) as % of the initial concentration.

| Microalgae | LI = 50 $\mu\text{mol m}^{-2} \text{s}^{-1}$ | LI = 100 $\mu\text{mol m}^{-2} \text{s}^{-1}$ | LI = 200 $\mu\text{mol m}^{-2} \text{s}^{-1}$ | LI = 400 $\mu\text{mol m}^{-2} \text{s}^{-1}$ |
| --- | --- | --- | --- | --- |
|  | % of initial |  |  |  |
| CCALA 242 | 17.6 | 8.1 | 15.1 | 16.2 |
| CCALA 243 | 92.5 | 95.5 | nd | nd |
| CCALA 244 | 85.5 | 84.6 | 67.6 | 41.4 |
| CCALA 453 | 10.9 | 10.6 | 12.9 | 10.3 |
| CCALA 455 | 17.1 | 10.4 | 6.0 | 6.5 |
| CCALA 456 | 0 | 0 | 0 | 0 |
| CCALA 463 | 0 | 0 | 0 | 0 |
| CCALA 464 | 29.2 | 10.5 | 7.7 | 0 |
| CCALA 467 | 9.8 | 12.5 | 9.3 | 6.3 |
| CCALA 688 | 11.5 | 5.2 | 5.1 | 0.3 |

In most cases, the highest amount of total fatty acids (TFA) was found in the biomass cultured at optimal light intensity (400 µmol m^−2^ s^−1^) (Fig. 3). Only the values higher than 10% of TFA were considered and this was the case for either palmitic acid (PA; C16:0), stearic acid (SA; C18:0), oleic acid (OA; C18:1n), linolenic acid (LA; C18:2 n6), or α-linolenic acid (ALA; C18:3n3). All microalgae were rich in PA as it was found in the amount ranging between 18.5 and 43.2% of TFA. The higher light intensities of 200 and 400 µmol m^−2^ s^−1^ favoured the production of PA as the highest concentration was found in the biomass of *C. moewusii* CCALA 244 in the amount of 43.2 and 42.3 % TFA, respectively. In the biomass of *D. communis* CCALA 463, the concentration of PA did not decrease bellow 34% of TFA regardless of light intensity. The third microalgae with the highest PA content was *D. subspicatus* CCALA 467, where its content at optimal illumination corresponded to 38% of TFA.The second most abundant FA found in the biomass was OA ranging from 10.4 to 37.7% of TFA. The production of OA was also supported by higher light intensity as the highest amount of 37.7 of TFAwas found in the biomass of *S. obliquus* CCALA 456 cultivated at 400 µmol m^−2^ s^−1^. The FA found in the biomass at the lowest concentration was SA C18:0, which was present from 10.2 to 14.8% of TFA.

**Fig. 3.**
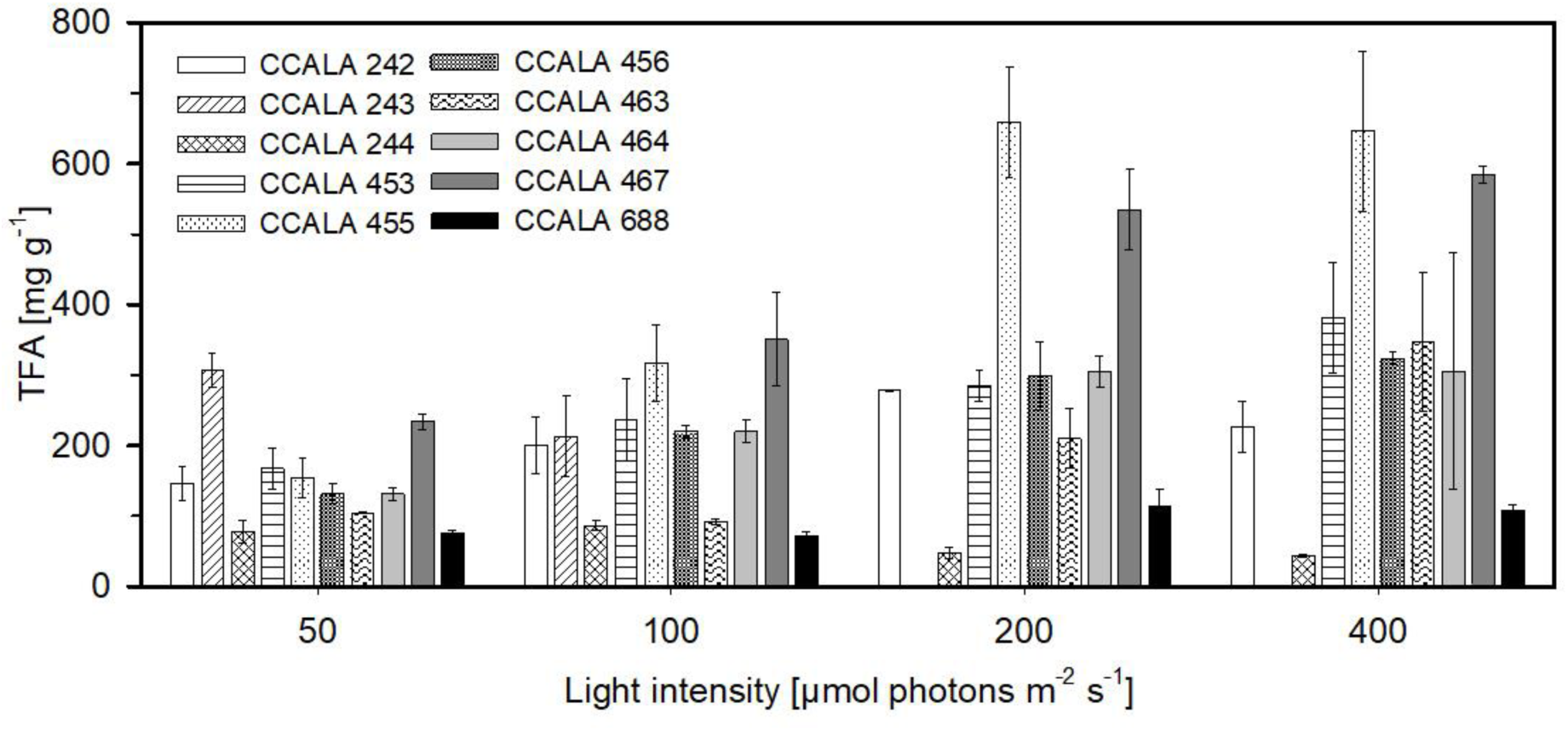
The total fatty acid content of selected microalgae (see legend of Fig. 1) analysed at the end of the trial (D10). The values are presented as a mean (n = 3) ± SD and those designated by the same letter did not differ significantly from each other.

**Fig. 4.**
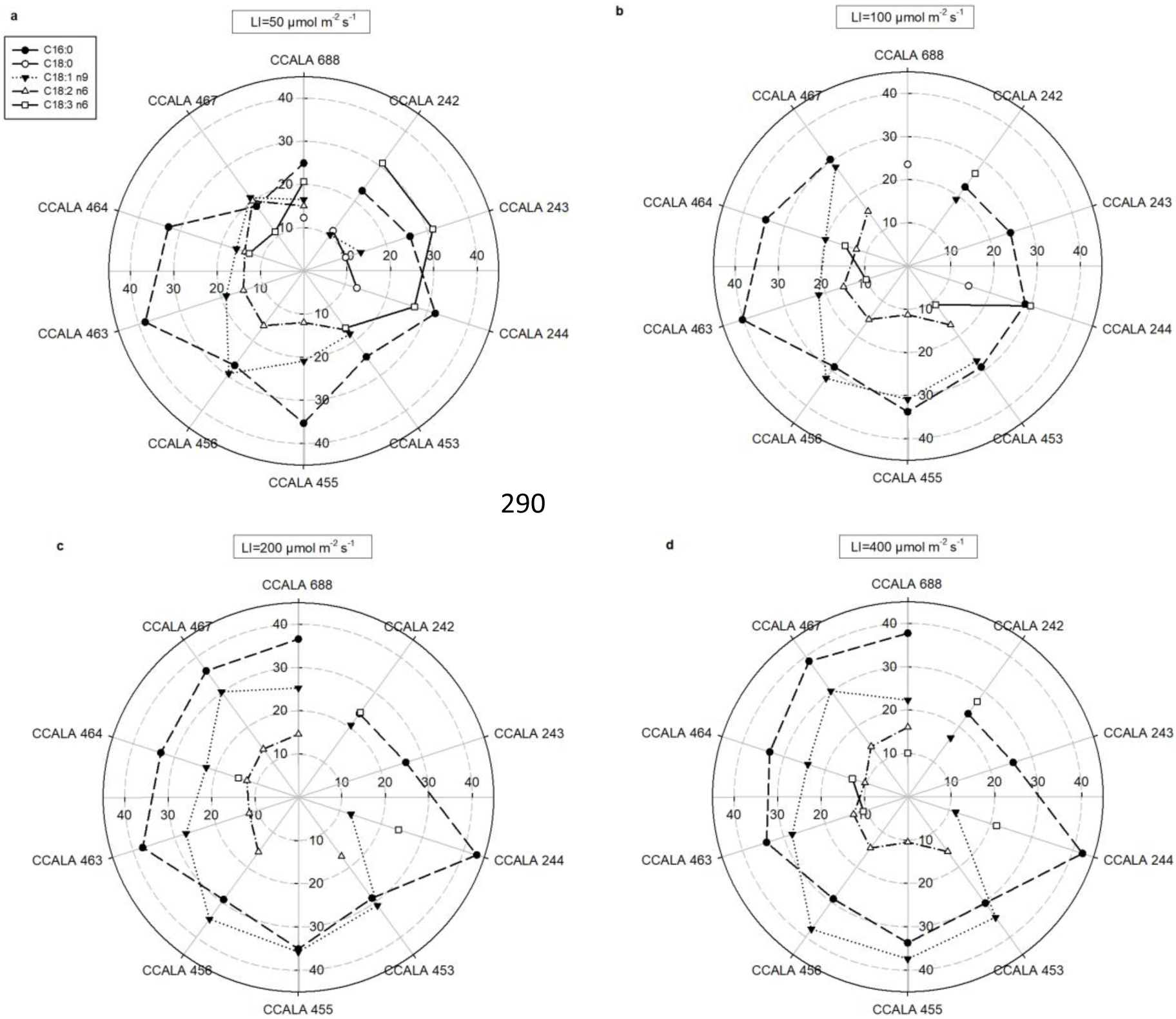
The concentration [%TFA] of five selected fatty acids (palmitic PA C16:0; stearic SA C18:0; oleic OA C18:1n9; linoleic LA C18:2 n6; α-linolenic C18:3 n3) analysed in the biomass samples of the microalgae cultures (list in the legend of Fig. 1) harvested at the end of the trial which were grown at different light intensities: a) 50 µmol m^−2^ s^−1^, b) 100 µmol m^−2^ s^−1^, c) 200 µmol m^−2^ s^−1^, d) 400 µmol _m_-2 _s_-1.

## Trial 2: The effect of unfavourable conditions on selected microalgae at optimum light intensity

In the second series of photoautotrophic experiments, the growth and FA production of *C. moewusii* CCALA 244, *D. communis* CCALA 463 and *D. subspicatus* CCALA 467 at a lower temperature of 25°C and a lower temperature of 25°C simultaneously in the presence of 16 g L^−1^ of NaCl were investigated when cultured in a small scale.

Both microalgae, *C. moewusii* CCALA 244 and *D. subspicatus* CCALA 467 grew well in the BG-11 medium (further as BG-11) even at a sub-optimal temperature of 25°C. The growthrate of *D. communis* CCALA 463 at suboptimal temperature in the BG-11 with 16 g L^−1^ NaCl (further as BG-11-NaCl) was affected and the highest culture density of 2.48g L^−1^ was measured on day 5 and then started to decrease. The highest densities were generally obtained when *D. subspicatus* CCLA 467 was cultured at sub-optimal temperature in both BG-11 without and with NaCl, reaching values of 6.48 g L^−1^ (D10) and 6.20 g L^−1^ (D7), respectively (Fig. 5).

**Fig. 5.**
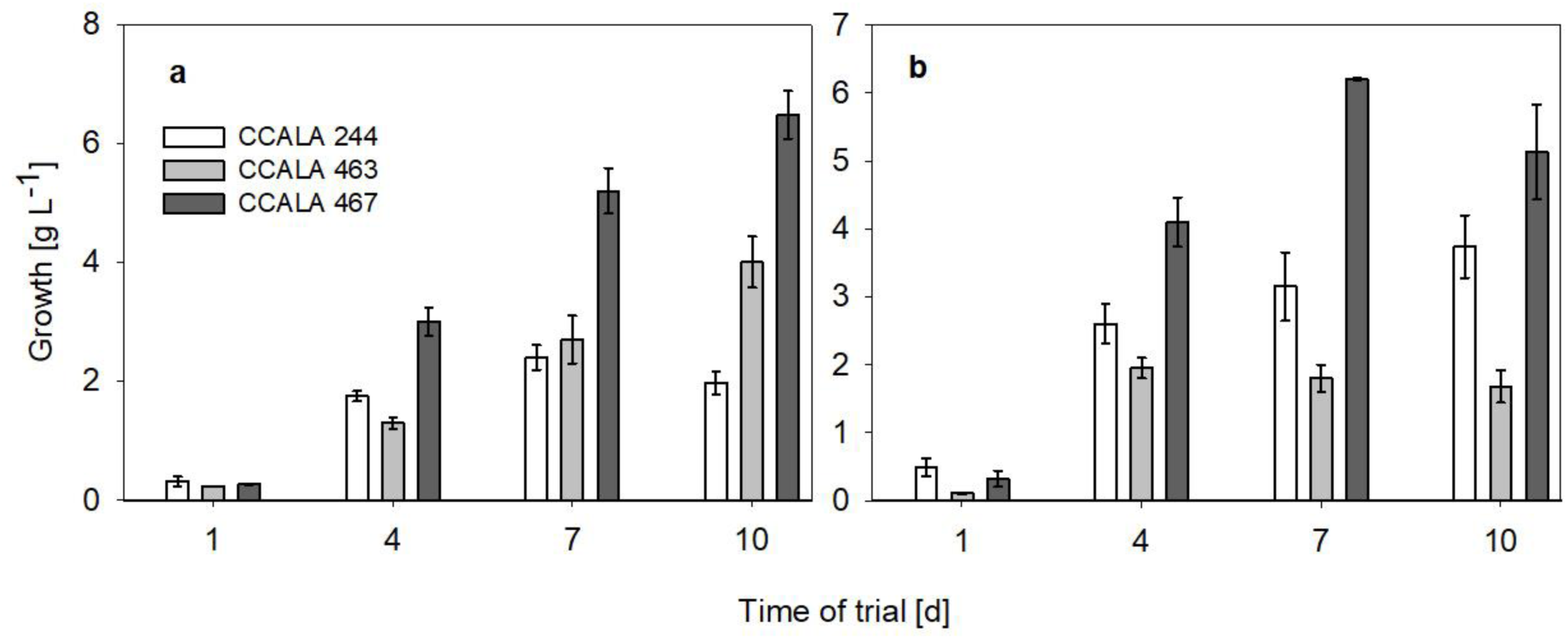
The growth of *Chlamydomonas moewusii* CCALA 244*, Desmodesmus communis* CCALA 463 and *Desmodesmus subspicatus* CCALA 467 at sub-optimal temperature of 25°C in (a) BG-11 and (b) in BG-11 supplemented with 16 g L^−1^ of NaCl (BG-11-NaCl). The values are presented as a mean (n = 3) ± SD and those designated by the same letter did not differ significantly from each other.

**Fig. 6.**
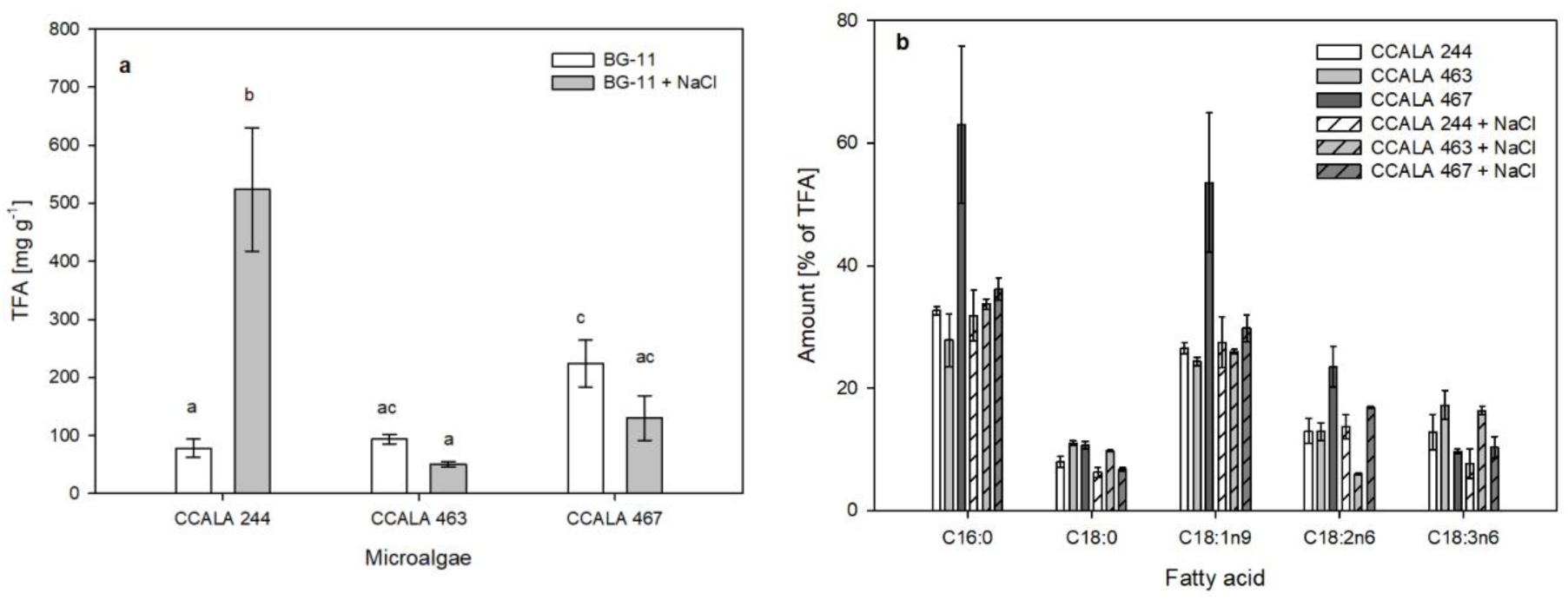
a) The total fatty acid (TFA) content [mg g^−1^] of three microalgae strains *C. moewusii* CCALA 244, *Desmodesmus communis* CCALA 463 and *Desmodesmus subspicatus* CCALA 467 and b) selected fatty acids (palmitic PA C16:0; stearic SA C18:0; oleic OA C18:1n9; linoleic LA C18:2 n6; α-linolenic C18:3 n3) analysed in the biomass samples of the microalgae cultures (list in the legend of Fig. 1) harvested at the end of the trial which were grown in the BG-11 and BG11-NaCl at 25 °C. The values are presented as a mean (n = 3) ± SD and those designated by the same letter did not differ significantly from each other.

The highest growth rates of µ=2.0 d^−1^ was calculated for *D. subspicatus* CCALA 467 regardless of the culture medium as the rates reached the same value (Table 3). The rapid growth also corresponds to the greatest nutrient loss measured at the end of the experiment (Table 4). Similar but lower growth rates regardless of the cultivation medium were also found for *C. moewusii* CCALA 244 – µ=0.2 d^−1^ and µ=0.3 d^−1^, respectively. The most significant difference in growth rates in the two different media used was found in the case of *D. communis* CCALA 463 as in BG-11-NaCl was approximately 3-fold higher reaching µ=1.0 d^−1^ compared with that reached in BG-11 (µ=0.3 d^−1^). In the fact the growth in BG-11 NaCl stagnated on D4 (Fig. 5b). The reluctance of *D. communis* of CCALA 463 to grow in the medium with added salt was also supported by the analysis of nitrate, where its amount measured at the end of the experiment in the medium was still 79.9±5.0% of the original concentration (Table 3).

**Table 3.** Growth rate [d^−1^] of selected microalgae cultivated at a sub-optimal temperature of 25°C in BG-11 medium and BG-11 medium with 16 g L^−1^ of NaCl.

| Microalgae | Growth rate [$d^{-1}$] | |
| --- | --- | --- |
| | BG-11 medium; $25^{\circ}C$ | BG-11 medium + $16 g L^{-1}$ NaCl; $25^{\circ}C$ |
| CCALA 244 | 0.3 | 0.2 |
| CCALA 463 | 0.3 | 1.0 |
| CCALA 467 | 2.0 | 2.0 |

**Table 4.** Nitrate (NO ^−^) concentration expressed as % of initial measured in the cultivation medium at the end of the trial.

| Microalgae | $NO_3^-$ concentration [% of initial] | |
| --- | --- | --- |
| | BG-11 medium; $25^{\circ}C$ | BG-11 medium + $16 g L^{-1}$ NaCl; $25^{\circ}C$ |
| CCALA 244 | $35.4 \pm 3.0$ | $53.1 \pm 5.8$ |
| CCALA 463 | $26.6 \pm 2.8$ | $79.9 \pm 5.0$ |
| CCALA 467 | $3.8 \pm 1.1$ | $16.7 \pm 0.0$ |

The amount of TFA in the biomass of *C. moewuaii* CCALA 244 and *D. sommunis* CCALA 463 at the end of the trial when grown in BG-11 were comparable reaching the values of abou 78 and 94 mg g^−1^ DW, respectively. On the other hand, a significant difference was found when cultivated in BG-11-NaCl as the amount of TFA in *C. moewusii* CCALA 244 biomass reached the value of 524 mg g^−1^ DW. The reverse effect of salt supplementation was observed in *D. communis* CCALA 463 as the TFA amount was not affected by the cultivation medium. On the other hand, the amount of PA, SA, OA as well as LA in *D. subspicatus* CCALA 467 biomass decreased significantly when cultivated in BG-11-NaCl. In general, the biomass of all three microalgae species selected for this trial, was richest in PA and OA as the amount ranged from 24.3 to 63.0 % of TFA.

## Trial 3: Validation of cultivation conditions for selected microalgae in larger-scale

In this trial, the growth and FA production of selected microalgae used in Trial 2 were validated at larger-scale cultivation in 30 litre annular column photobioreactor (AC-PBR).

The culture of *D. subspicatus* CCALA 467 grew well regardless of whether NaCl was added to the BG-11medium or not. The culture grew linearly throughout the trial. Initial faster growth was observed in BG-11-NaCl (Fig. 7). Stationary phase of growth was observed in both experiments from D15 as the culture became nutrient-limited (Table 6). The concentration of nitrate dropped to 8-9% of the initial concentration on D16. The maximum biomass density reached by *D. subspicatus* CCALA 467 in both media was hight and reached 5.2 g L^−1^ in BG-11 and 5.9 g L^−1^ when BG-11-NaCl was used.

**Fig. 7.**
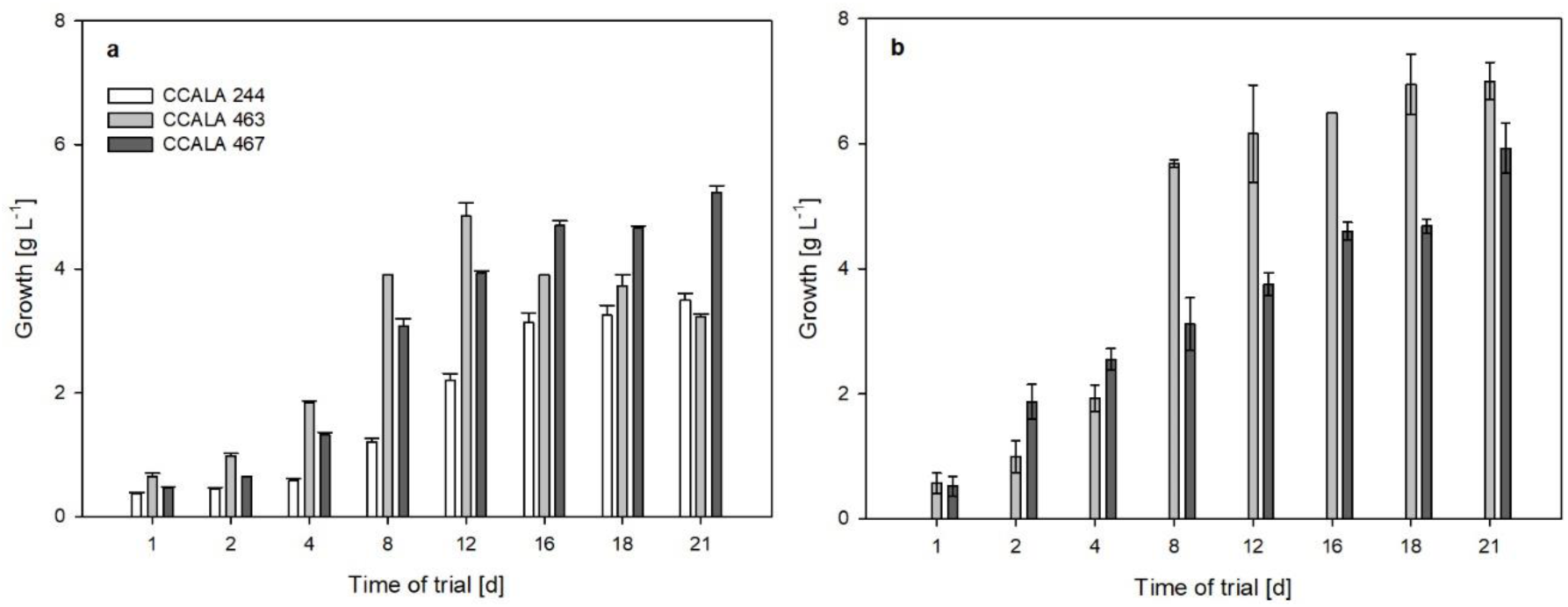
The growth of *Chlamydomonas moewusii* CCALA 244*, Desmodesmus communis* CCALA 463 and *Desmodesmus subspicatus* CCALA 467 at sub-optimal temperature of 25°C in (a) BG-11 medium and (b) in BG-11-NaCl in 30 L AC-PBR. The values are presented as a mean (n = 3) ± SD and those designated by the same letter did not differ significantly from each other.

Compared to the cultures grown in BG-11 (P_V_=0.24 g L^−1^ d^−1^, P_A_=0.18 g m^−2^ d^−1^, respectively) slightly higher volumetric as well as areal productivities were calculated for the *D. subspicatus* CCALA 467 biomass grown in BG-11-NaCl reaching the values of P_V_=0.27 g L^−1^ d^−1^ and P_A_=0.20 g m^−2^ d^−1^. The volumetric and areal productivities reached in BG-11 medium were P_V_=0.24 g L^−1^ d^−1^ and P_A_=0.18 g m^−2^ d^−1^, respectively (Table 5).

**Table 5.** The volumetric (P_V_; g L^−1^ d^−1^) and areal productivities (P_A_; g m^−2^ d^−1^) of selected microalgae cultivated at sub-optimal temperature of 25°C in BG-11 medium and in BG-11 medium with 16 g L^−1^ of NaCl in 30 L AC-PBR.

| Microalgae | $P_V [\text{g L}^{-1} \text{ d}^{-1}]$ | | $P_A [\text{g m}^{-2} \text{ d}^{-1}]$ | |
| --- | --- | --- | --- | --- |
|  | BG-11 medium | BG-11 medium + NaCl | BG-11 medium | BG-11 medium + NaCl |
| CCALA 244 | 0.14 | 0 | 0.11 | 0 |
| CCALA 463 | 0.13 | 0.32 | 0.10 | 0.24 |
| CCALA 467 | 0.24 | 0.27 | 0.18 | 0.20 |

**Table 6.**
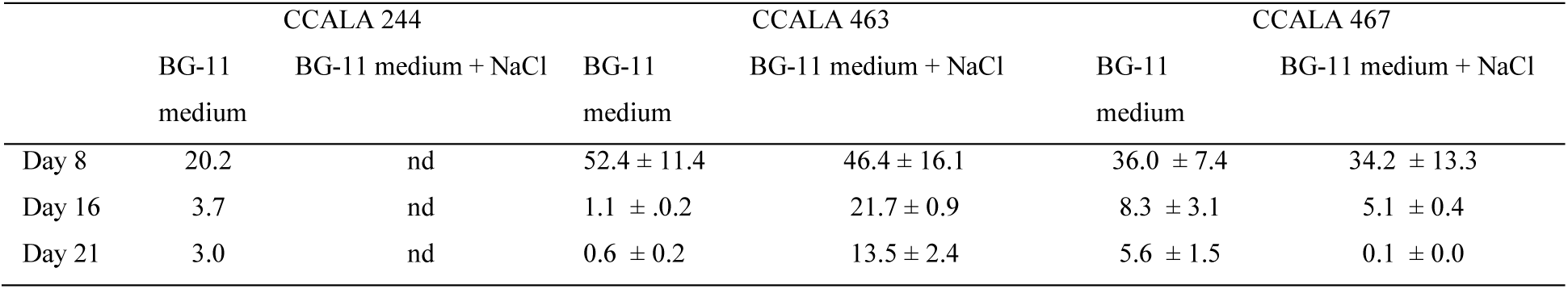
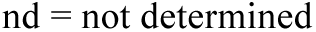
Nitrate (NO_3_^−^) concentration expressed as % of the initial value measured in the cultivation medium at the end of the trial in 30 L AC-PBR.

|  | CCALA 244 |  | CCALA 463 |  | CCALA 467 |  |
| --- | --- | --- | --- | --- | --- | --- |
|  | BG-11 medium | BG-11 medium + NaCl | BG-11 medium | BG-11 medium + NaCl | BG-11 medium | BG-11 medium + NaCl |
| Day 8 | 20.2 | nd | $52.4 \pm 11.4$ | $46.4 \pm 16.1$ | $36.0 \pm 7.4$ | $34.2 \pm 13.3$ |
| Day 16 | 3.7 | nd | $1.1 \pm 0.2$ | $21.7 \pm 0.9$ | $8.3 \pm 3.1$ | $5.1 \pm 0.4$ |
| Day 21 | 3.0 | nd | $0.6 \pm 0.2$ | $13.5 \pm 2.4$ | $5.6 \pm 1.5$ | $0.1 \pm 0.0$ |

nd = not determined

The highest content of TFA was observed in *D. subspicatus* CCALA 467 strain on D11 reaching the value of 93.7 ± 1.6 µg mg^−1^ when cultivated in BG-11 medium and 92.7±1.2 µg mg^−1^ in BG-11-NaCl medium (Fig. 8a). However, the biomass in both cases contained significant amounts of only PA; OA was present in smaller amounts (maximum reached app. 10% of TFA). In general, a higher concentration of PA was present in the biomass cultivated in BG-11 as the concentration ranged between 21.9 ± 2.6 and 27.3 ± 5.9% of TFA while in the biomass cultivated in BG-11 medium with salt addition, the highest amount of PA reached only 25.0 ± 0.1% of TFA on D21 (Fig. 8b).

**Fig. 8.**
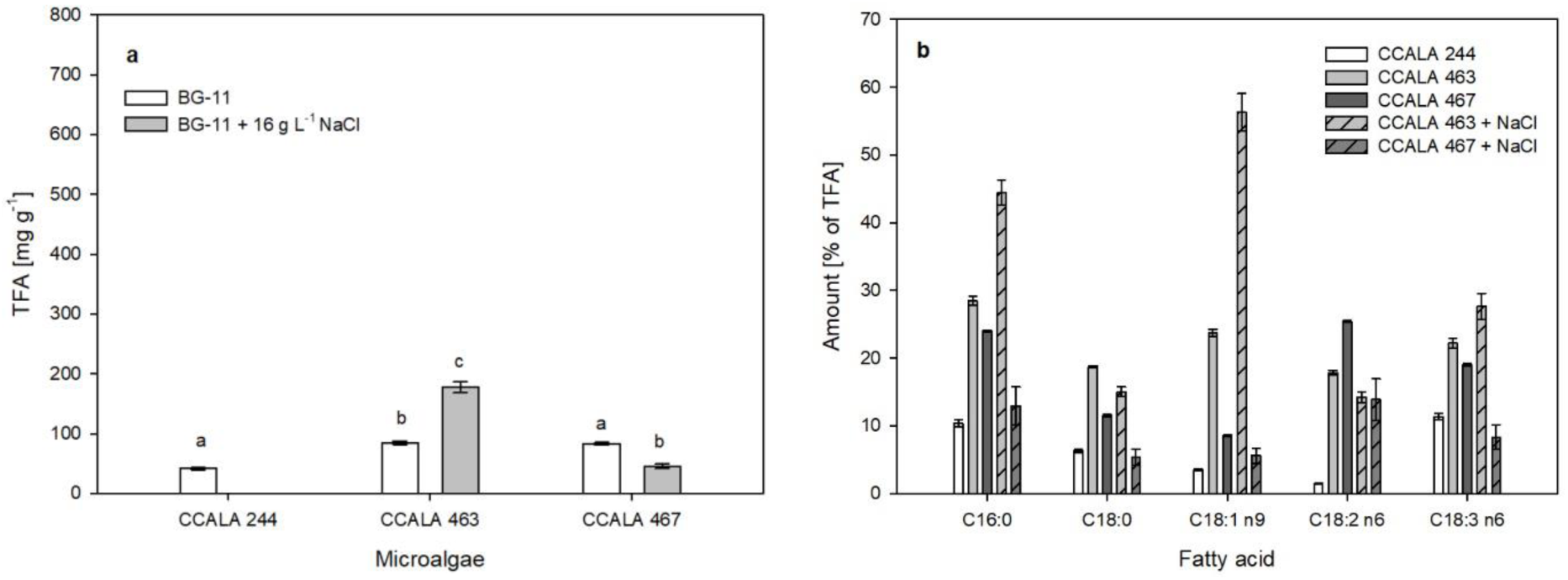
a) The total fatty acid (TFA) content [mg g^−1^] of three microalgae strains *C. moewusii* CCALA 244, *Desmodesmus communis* CCALA 463 and *Desmodesmus subspicatus* CCALA 467 cultivated in 30 L AC-PBR and b) the content of selected fatty acids (palmitic PA C16:0; stearic SA C18:0; oleic OA C18:1n9; linoleic LA C18:2n6; α-linolenic C18:3n3) analysed in the biomass samples of the microalgae cultures (list in the legend of Fig. 1) harvested at the end of the trial which were grown in the BG-11 and BG-11-NaCl at 25 °C. The values are presented as a mean (n = 3) ± SD and those designated by the same letter did not differ significantly from each other.

A different trend of biomass growth was observed during the cultivation of *D. communis* CCALA 463 strain. The initial two-day lag phase was observed during the cultivation in the BG-11-NaCl medium, then the culture grew linearly until D11 when the culture probably became light-limited as there was still a noticeable amount of nitrate at the end of the experiment (Table 6) and a further increase in the biomass density was moderate. The same linear growth but without lag phase was evident when the culture was grown in the medium without salt addition. The biomass density increased linearly as in the previous case until D11 and then the density started to decrease due to nutrient limitation (Fig. 7). The volumetric and areal productivities differed when cultured in the two media – BG-11 and in BG-11-NaCl. The volumetric productivity of biomass cultivated in BG-11 was P_V_=0.13 g L^−1^ d^−1^ while that in BG-11-NaCl was P_V_=0.32 g L^−1^ d^−1^. This corresponded to areal producctivities of P_A_=0.10 g m^−2^ d^−1^ and P_A_=0.24 g m^−2^ d^−1^ (Table 4). The TFA content increased linearly throughout the cultivation of *D. communis* CCALA 463 in the AC-PBR grown in BG-11. From an initial 42.0±5.4 µg mg^−1^, the content of TFA doubled to 84.4±2.8 µg mg^−1^. The concentration of PA decreased during the experiment while the concentration of OA increased. From D11 the concentration of PA and OA did not change significantly and at the end of the trial the concentration was 28.5±0.7 and 23.7±0.6 % of TFA, respectively. The TFA content in the biomass of *D. communis* CCALA 463 cultivated in BG-11-NaCl increased during the growth reaching the maximum of 217.9±10.3 µg mg^−1^ DW, which is 2.5 times higher than the maximum found in biomass cultivated without salt supplementation (Fig. 8a). The initial concentration of PA was 24.9±0.8% of TFA and decreased to 20.0±0.1% of TFA on D7 and then gradually increased again until it reached a maximum of 25.0 ±0.3% of TFA on D21. The amount of PA in the biomass cultured in both media was almost identical but the amount of OA differed significantly. The highest concentration of OA was found in the biomass cultured in BG-11 and reached the maximum of 23.9±2.3% of TFA on D16 while the highest concentration found in the biomass cultured in BG-11-NaCl reached the maximum value of 32.3±0.1% of TFA on D16 (Fig. 8b). The culture of *C. moewusii* CCALA 244 strain grew well in BG-11 in BG-11-NaCl the culture did not grow, faded and the experiment failed after 3 days (data not shown). The increase in the biomass density was linear, reaching the value of 3.3±0.2 g L^−1^. The concentration of nitrate decreased linearly until the concentration on the last day was still approximately 4% of the original concentration (Table 6). The volumetric productivity of the biomass grown in AC-PBR was P_V_=0.14 g L^−1^ d^−1^ corresponding to the areal productivity of P_A_=0.11 g m^−2^ d^−1^. The TFA content ranged between 41.5±1.8 and 71.4±1.9 mg g^−1^ DW (Fig. 8a). The initial concentration of PA as well as OA dropped during the first two days to 21.4±0.4 and 10.2±0.4% of TFA, respectively. Thereafter, the concentrations of both FAs increased linearly until they reached the maximum values of 25.1±0.2 and 8.4±0.0% of TFA on the last day of the trial (Fig. 8b).

## Trial 4: Heterotrophic experiments

In Trial 4, the accumulation of biomass and maximum growth rate of eight heterotrophically growing microalgae was determined (Fig. 9). The strains split into two groups. The first group (*C. moewusii* CCALA 244, S*. obliquus* CCALA 455, *D. communis* CCALA 463 and *D. communis* CCALA 464) grew faster and reached higher maximum biomass after about 50 to 60 hours of cultivation. *C. moewusii* CCALA 244 grew fastest from all the strains although its final maximum biomass was among average of the first group.

**Fig. 9.**
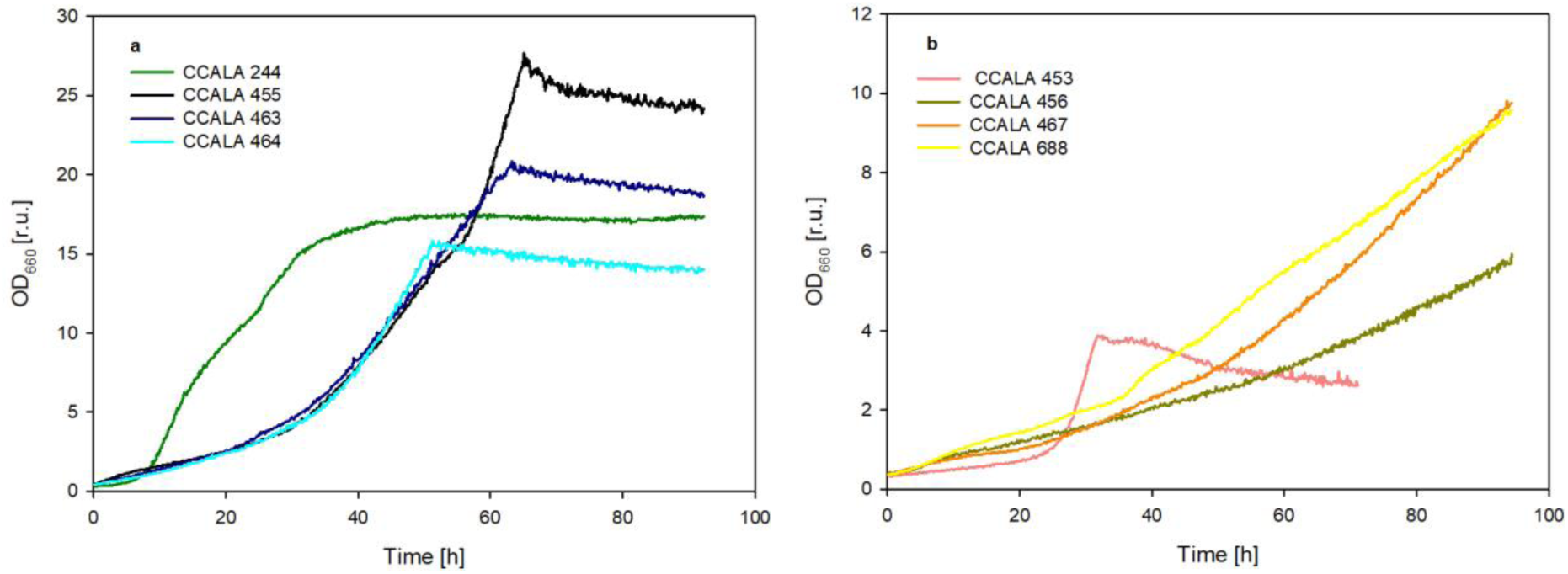
Growth of selected microalgae (*Scenedesmus obliquus* strains CCALA 453, CCALA 455 and CCALA 456, *Chlamydomonas moewusii* strain CCALA 244, *Desmodesmus communis* strains CCALA 463 and CCALA 464, *Scenedesmus subspicatus* CCALA 688, *Desmodesmus subspicatus* CCALA 467) followed during heterotrophic growth in the presence of glucose for 4 days. Representative growth curves from three independent experiments are presented.

The second group (*S. obliquus* CCALA 453, *S. obliquus* CCALA 456, *D. subspicatus* CCALA 688 and *D. subspicatus* CCALA 467) grew about twice slower and reached lower maximum biomass at the end of the experiment. The maximum growth rates (Fig. 10) reflected the growth patterns and were generally bigger than these of autotrophically grown cultures (see comparison in Table 7). When comparing growth rates suitable light intensity of all microalgae was determined based on the accumulation of the biomass and therefore the calculated maximum growth rate.

**Fig. 10.**
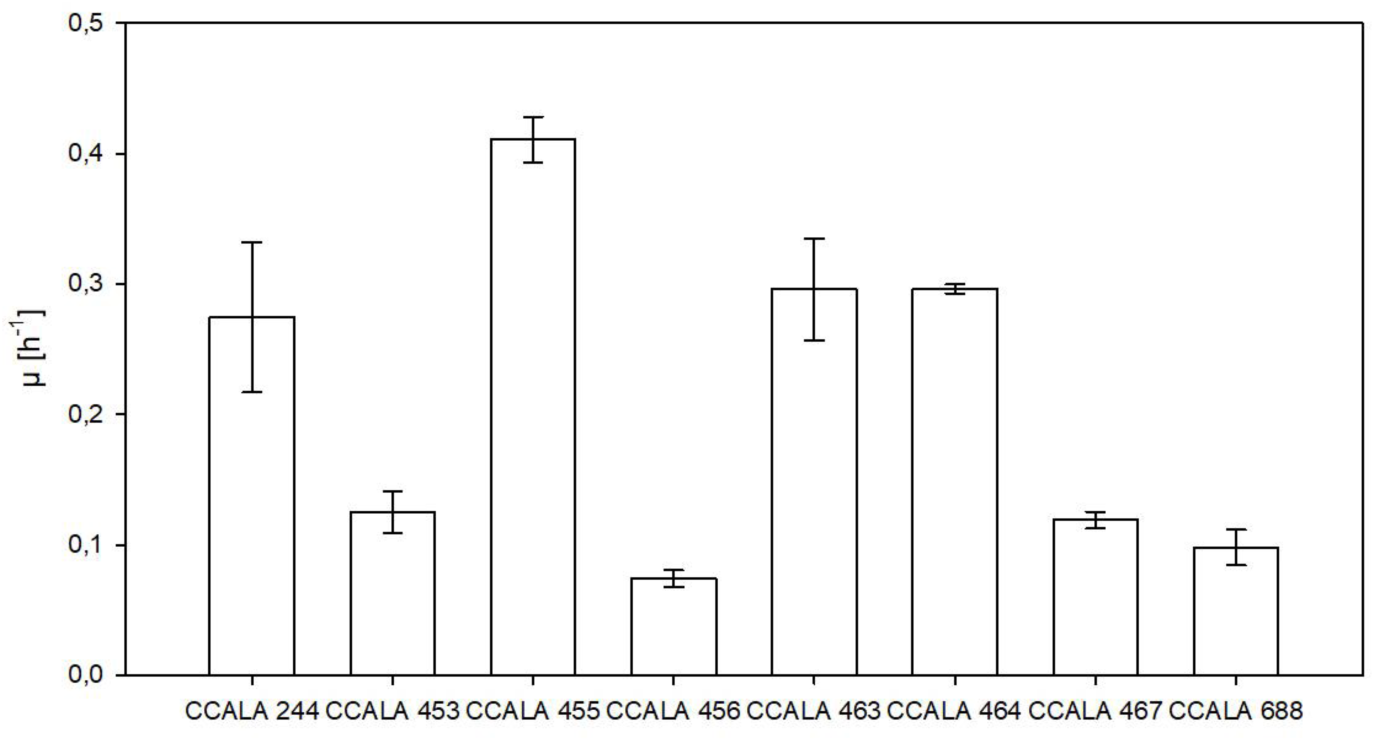
Maximum growth rates of selected microalgae strains (see legend of Fig. 1) grown heterotrophically on 1% glucose at 30 °C. *C. moewusii* CCALA 242 and 243 did not grow heterotrophically. The values are presented as a mean (n = 3) ± SD and those designated by the same letter did not differ significantly from each other.

**Table 7.** Ratio of growth rates of selected microalgae strains (see legend of Fig. 1) grown heterotrophically on 1% glucose at 30 °C compared to the maximum growth rate reached in autotrophic experiments. *C. moewusii* CCALA 242 and 243 did not grow heterotrophically.

| Microalgae strain | heterotrophy vs autotrophy |
| --- | --- |
| CCALA 242 | NA |
| CCALA 243 | NA |
| CCALA 244 | 22.43 |
| CCALA 453 | 6.05 |
| CCALA 455 | 20.41 |
| CCALA 456 | 3.22 |
| CCALA 463 | 5.69 |
| CCALA 464 | 15.26 |
| CCALA 467 | 5.67 |
| CCALA 688 | 4.21 |

In the heterotrophically grown cultures, the TFA in all biomass samples was always significantly lower as compared autotrophically grown biomass. The most significant difference in TFA content was determined for *S. subspicatus* CCALA 688 biomass, with heterotrophically grown biomass containing 30 times less TFA than autotrophically grown biomass. (compare Fig. 3 and Fig. 11a). Of the 5 FAs that were making the highest proportion in autotrophic cultures (palmitic acid PA C16:0; stearic acid SA C18:0; oleic acid OA C18:1n9c; linolenic acid LA C18:2n6; and α-linolenic acid C18:3n3) the most abundant FA was PA ranging from 46.85 % of TFA in *C. moewusii* CCALA 244 to 5.97 % in *D. communis* CCALA 464 (Fig. 11b). Stearic acid was very low in majority of the cultures reaching the values between 0.5-0.7 % of TFA, only in *C. moewusii* CCALA 244 (15.89 % of TFA) and in *S. obliquus* CCALA 453 (4.49 % of TFA). Similarly, OA and LA was detected in higher amount only in *C. moewusii* CCALA 244 (7.17 % of OA, 1.82 % LA) and *S. obliquus* CCALA 453 (17.53 % of OA, 5.91 % LA).

**Fig. 11.**
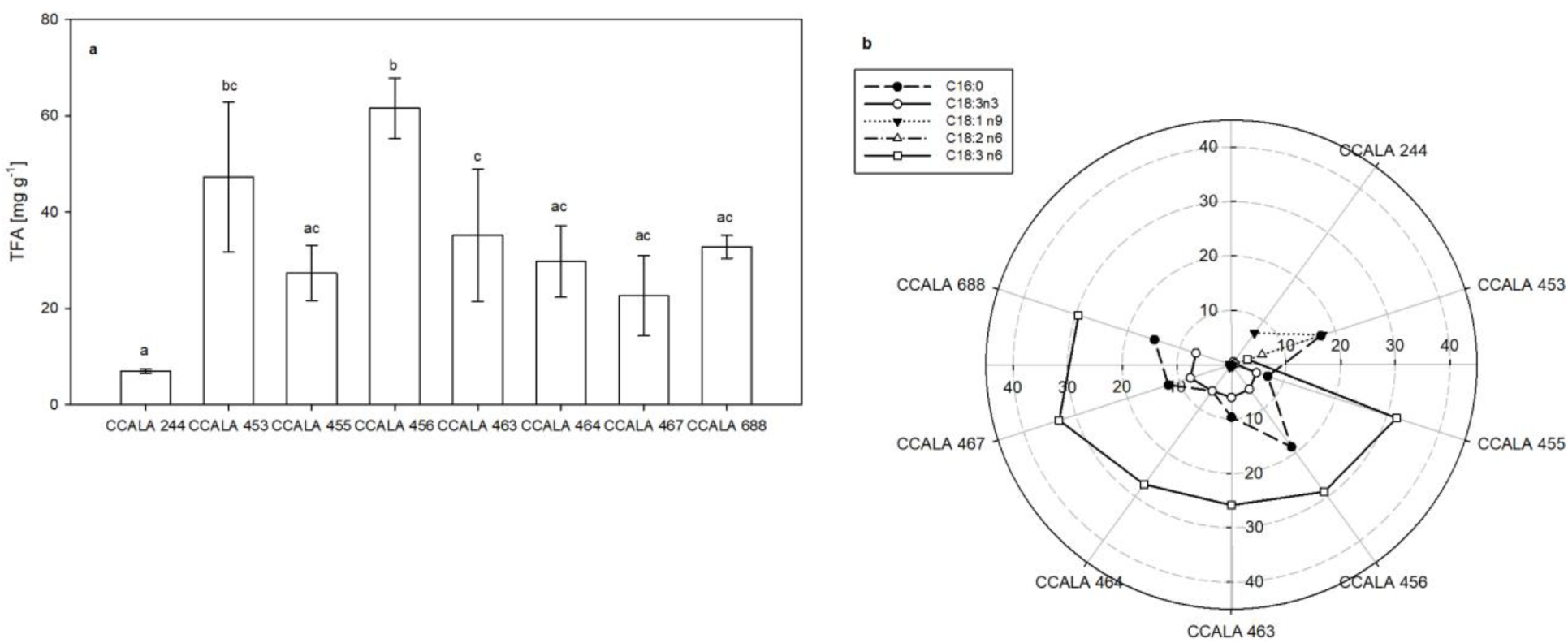
Fatty acid production of selected microalgae strains (see legend of Fig. 1) grown heterotrophically on 1% glucose at 30 °C a) the total fatty acid (TFA) content [mg g^−1^] and b) the concentration [%TFA] of five selected fatty acids (palmitic PA C16:0; stearic SA C18:0; oleic OA C18:1n9; linoleic LA C18:2 n6; α-linolenic C18:3 n3) analysed in the biomass samples harvested at the end of the trial. The values are presented as a mean (n = 3) ± SD and those designated by the same letter did not differ significantly from each other.

In contrast, γ-linolenic acid C18:3n6 was one of the most abundant FAs in the heterotrophically grown algae ranging from 3.10 % in *S. obliquus* CCALA 453 to 33.14 % in *D. subspicatus* CCALA 467. Interestingly, it was absent from *C. moewusii* CCALA 244.

## Discussion

Algae adapt both their growth rates and the fatty acids production to growth conditions, with two critical factors under non-stress conditions being temperature and light [14,42,43]. Here, we selected a set of algae that naturally produce oils similar in composition to palm oil, and we analyzed the effect of varying light intensity on their growth and fatty acid production and composition while maintaining temperature. Two of the three *C. moewusii* strains (CCALA 242 and 243) preferred low light intensities, with *C. moewusii* CCALA 243 growing best at the lowest light intensity and *C. moewusii* CCALA 242 preferring the low light intensity of 100 µmol m^−2^ s^−1^. Such a low light preference was previously shown for *C. eugametos* (synonymous with *C. moewusii*) [44], which was routinely grown at around 100 µmol m^−2^ s^−1^ and 150 µmol m^−2^ s^−1^ [45]. In contrast, the most studied member of the genus Chlamydomonas, *Chlamydomonas reinhardtii*, can grow at different light intensities from 6 to 250 µmol m^−2^ s^−1^ [46] and was not saturated even at 500 µmol m^−2^ s^−1^ [47]. The growth of *Scenedesmus* and *Desmodesmus* strains improved with increasing light intensity. The genera *Scenedesmus* and *Desmodesmus* include numerous widely studied biotechnologically relevant species. Their response to light appears to be dependent on both the specific species/strain and the growth conditions. For flask cultures of various *Scenedesmus* species, the optimal growth light intensity was 81 µmol m^−2^ s^−1^ [48]. Cultures of *Scenedesmus obliquus* grown in flatbed reactors achieved maximum growth rates at around 150 µmol m^−2^ s^−1^, although they were able to tolerate and adapt to higher light [49]. *Scenedesmus* sp. cultured in tubular reactors saturated at 250 µmol m^−2^ s^−1^ and maintained similar biomass yields at 400 µmol m^−2^ s^−1^ [50]. *Scenedesmus* quadricauda grown in a tubular photobioreactor preferred a higher light intensity of 500 µmol m^−2^ s^−1^ [51].

Stress conditions are often used to increase the production of lipids and improve their productivity. One of the most commonly used stress factors is nitrogen limitation, which leads to a reduction in growth but promotes lipid accumulation [16]. Another stress factor that is easy to manage is salinity. Under mild salt stress, lipid content increases [26–28], while higher concentrations can have a detrimental effect. In our experiments, the effect of NaCl administration (approx. 280 mM) varied depending on the species. The salinity improved the total fatty acid production only in *C. moewusii* CCALA 244, although the amount of PA, SA, OA was comparable between the two condition. A similar effect of salinity on lipid production was shown for *Chromochloris zofingiensis* [52], a close relative of *Chlamydomonas reinhardtii* [53]. In *C. zofingiensis*, the application of salt led to an approximately threefold increase in triacylglycerol content and an approximately fourfold increase in asthaxanthin content, while biomass productivity decreased by only 20%. [52]

Apart from the detrimental effects of increased salinity on the growth and TFA content of *D. communis* CCALA 463, cultivation of *D. subspicatus* CCALA 467 in NaCl-enriched medium resulted in a reduction in total fatty acid production as well as particularly affected the production of PA, SA, OA and LA. This higher sensitivity to salinity is consistent with the demonstrated sensitivity of the genera *Desmodesmus* and *Scenedesmus* to salt concentrations above 100 mM [54] or above 160 mM [55]. The differential effect of salinity on the two related species is consistent with previous reports on the effect of salinity on *Scenedesmus quadricauda* and *Scenedesmus dimorphus*, where each species responded differently. In *Scenedesmus quadricauda*, both biomass and lipid content were significantly increased at 160 mM NaCl. In contrast, *Scenedesmus dimorphus* [55] showed no difference in biomass productivity or lipid content at the same NaCl concentration, although both decreased at higher salinity (320 mM NaCl) [55]. Other species of the genus *Scenedesmus* have proven to be more resistant. *Scenedesmus* sp. BHU1 showed an approximately threefold decrease in biomass accumulation but a fourfold improvement in lipid and carbohydrate content in 400 mM NaCl [56].

In large-scale cultivation, the growth rates of all three species (*C. moewusii* CCALA 244, *D. subspicatus* CCALA 467 and *D. communis* CCALA 463) were comparable to small-scale growth under unstressed conditions and the biomass productivity of *D. communis* CCALA 463 and *D. subspicatus* CCALA 467 was similar to that of *S. almariensis* [57], although it was lower than that of *S. almariensis* at higher light intensities [58,59] and *S. obliquus* [60].

Many algae species are capable of heterotrophic growth. Such growth can be faster and the final cell densities can be higher than in autotrophy, as the growth density is not limited by light [61]. This is advantageous for further processing of the algal biomass and can significantly reduce production costs despite the addition of an organic carbon source. Of the ten algae species in this study, eight were able to grow heterotrophically. All the strains grew significantly better in heterotrophy. In contrast, the maximum biomass achieved by the two treatments was not comparable. In heterotrophy, it varied between 1 and 3 g/l and the highest value was 4.8 g/l in *S. obliquus* CCALA 455, which is comparable to the lowest light intensity, but not to the higher ones. This is surprising, as heterotrophic biomass productivities are generally high, ranging from 20 g/l to 286 g/l, with higher growth rates typical for different species of the genus *Chlorella* sp. and the highest biomass achieved by *Scenedesmus acuminatus* [62]. With similar overall productivity of FAs between autotrophy and heterotrophy, heterotrophy seems to be suitable for FA production, particularly due to the improved growth rates.

## Acknowledgments

The authors thank Ms. Soňa Pekařová, Ms. Kamila Ondrejmišková, Ms. Klára Kochtová and Mr. Martin Lukeš for technical assistance and Dr. Richard Lhotský for administrative support.

## Author contributions

Conceptualization, K.Š., K.B.; methodology, K.Š., K.B., J.A.C.M.; investigation, K.Š., K.B., J.A.C.M.; data curation, K.Š., K.B., J.A.C.M., J.M.; writing—original draft preparation, K.Š., K.B., J.A.C.M., J.M.; writing—review and editing, K.Š., K.B., J.A.C.M., J.M.; visualization, K.Š., K.B., J.M.; supervision, K.Š., K.B., J.M.; funding acquisition, K.Š., K.B. All authors have read and agreed to the published version of the manuscript.

## Data availability

The datasets presented in this study are available from the corresponding authors upon reasonable request. The data are not publicly available without the permission of all co-authors.

